# Spatially resolved quantification of oxygen consumption rate in ex vivo lymph node slices

**DOI:** 10.1101/2024.01.03.573955

**Authors:** Parastoo Anbaei, Marissa G. Stevens, Alexander G. Ball, Timothy N.J. Bullock, Rebecca R. Pompano

## Abstract

Cellular metabolism has been closely linked to activation state in cells of the immune system, and the oxygen consumption rate (OCR) in particular serves as a valuable metric for assessing metabolic activity. Several oxygen sensing assays have been reported for cells in standard culture conditions. However, none have provided a spatially resolved, optical measurement of local oxygen consumption in intact tissue samples, making it challenging to understand regional dynamics of consumption. Therefore, here we established a system to monitor the rates of oxygen consumption in ex vivo tissue slices, using murine lymphoid tissue as a case study. By integrating an optical oxygen sensor into a sealed perfusion chamber and incorporating appropriate correction for photobleaching of the sensor and of tissue autofluorescence, we were able to visualize and quantify rates of oxygen consumption in tissue. This method revealed for the first time that the rate of oxygen consumption in naïve lymphoid tissue was higher in the T cell region compared to the B cell and cortical regions. To validate the method, we measured OCR in the T cell regions of naïve lymph node slices using the optical assay and estimated the consumption rate per cell. The predictions from the optical assay were similar to reported values and were not significantly different from those of the Seahorse metabolic assay, a gold standard method for measuring OCR in cell suspensions. Finally, we used this method to quantify the rate of onset of tissue hypoxia for lymph node slices cultured in a sealed chamber and showed that continuous perfusion was sufficient to maintain oxygenation. In summary, this work establishes a method to monitor oxygen consumption with regional resolution in intact tissue explants, suitable for future use to compare tissue culture conditions and responses to stimulation.

**TOC image:** 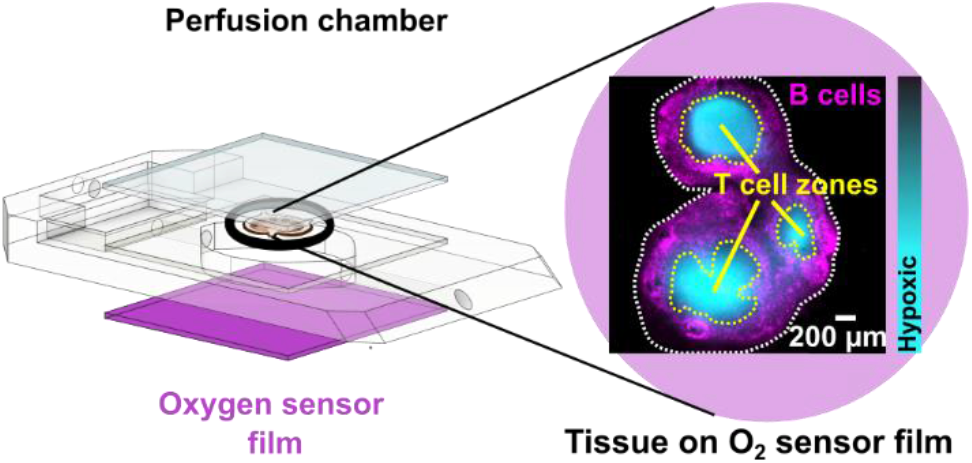

## 2 INTRODUCTION

Measurement of oxygen metabolism plays a significant role in determining the metabolic state and functioning of tissues, especially in the context of immunity where cellular activation is closely associated with metabolic state. The impact of hypoxia on cell state has been studied for decades, revealing, for example, the role of hypoxia-inducible factor-1 (HIF-1α) in human cancers and infectious disease and the potential to target this pathway for immunotherapy.^1–3^ Measurement of regions of depleted oxygen, made by using hypoxia-sensitive fluorescent probes injected in vivo, has yielded insights such as the role of hypoxia in controlling B cell function in germinal centers.^4,5^ Meanwhile, direct measurement of oxygen consumption rate (OCR) by the Seahorse Mito-Stress Test assay provided widely used functional readouts that were central to understanding T cell and B cell metabolic states,^6^ and were recently extended to use with small intact tissue slices.^7^ However, no regional spatial information is provided using this method, as it is based on bulk measurement in a microchamber containing cells or organoids in culture.

Analysis of oxygen consumption rate in live, organized tissues potentially offers a more realistic understanding of oxygen dynamics than in cell culture settings.^8^ In vitro analyses of isolated cells might not accurately represent in vivo conditions due to the absence of local tissue architecture and related intercellular interactions, and the loss of the structural and regulatory support provided by extracellular matrix. One route to retaining this organization is to use ex vivo tissue slices. Although lower throughput than cell culture studies, ex vivo models enable researchers to observe dynamic cellular behaviors in the context of regional variations in the tissue. Furthermore, like traditional cell cultures, ex vivo tissue slices can be monitored over time in response to controlled changes in culture conditions.^8^

Several methods are available for measuring oxygen consumption rates in tissue, but none offer spatially resolved analysis of tissue substructures coupled with quantification of oxygen consumption rate. Methods such as positron emission tomography coupled with radiolabeled tracers,^9,10^ electron paramagnetic resonance oximetry,^11^ magnetic resonance imaging^12–14^, and near-infrared monitoring,^15^ are suitable for in vivo imaging but have limited spatial resolution (usually on the scale of mm or hundreds of microns), limited quantification of concentration, and high cost.^16^ Other methods utilize electrode-based sensors to provide structural and dynamic information even deep within tissue, but the probes are invasive and usually provide local data from a few specific points rather than an image.^17–21^ In contrast, optical methods based on visible light may not be suitable for deep in vivo locations due to light scattering, but they can easily provide regional or cellular spatial resolution, high temporal resolution depending on the kinetics of the selected dye, and an image if coupled with microscopy.^16,22,23^ A common strategy is to make use of luminescent (fluorescent or phosphorescent) dyes that are quenched by oxygen.^16^ Though often used for point-based sensors, some luminescent dyes have been coupled with imaging to great effect to monitor oxygen levels in 3D cultures^24^, as well as in mouse and rat brain, spleen, bone marrow, and retina tissue.^5,25–31^ However, accurately quantifying regional variations in oxygen consumption rate (OCR) within live, spatially organized tissue structures still poses a significant challenge due to tissue heterogeneity and the need to co-register oxygen signal with other markers of tissue geography while measuring dynamic metabolic events, and the need to maintain tissue viability during imaging.

Therefore, here we sought to design a system to visualize and quantify the rate of oxygen consumption in live tissue slices ex vivo with regional spatial resolution. We selected murine lymph node slices as a case study, as immunometabolism is an exciting area of interest for immunotherapy development. Information on the distribution of oxygen consumption in the lymph node is limited so far to a few local measurements made by implanted electrodes.^17,19^ According to dyes sensitive to hypoxic conditions, germinal centers in reactive lymph nodes may be more hypoxic than other areas of the lymph node,^4,5^ but no information is available on relative rates of oxygen consumption in various regions of the naïve lymph node. We have previously shown that ex vivo lymph node slices retain the spatial organization of the organ, as well as immune and metabolic activity in acute cultures.^32,33^ Therefore, here we took advantage of luminescent oxygen-sensing films in combination with live ex vivo slice culture to develop an imaging-based method to map out regional oxygen consumption rate in lymph node tissue ex vivo and to test the hypothesis that different regions consumed oxygen at different rates in naïve lymph node tissue.

## 3 EXPERIMENTAL

### 3.1 Preparation of oxygen-sensing glass slides

22×22 mm glass coverslips (Ted Pella, USA) were cleaned with 75% ethanol and then silanized by immersion in a solution of m,p-ethyl phenethyl trimethoxy silane (Gelest, USA) in ethanol (1:10 v/v ratio of silane: 200 proof ethanol) for 30 minutes, to decrease the polarity of their surface.^34^ After silanization, the slides were rinsed with EtOH and dried with nitrogen before spin coating. Oxygen-sensing glass slides were fabricated using a protocol developed by Lockett laboratory at the University of North Carolina.^24^ A cocktail of 1 mM Palladium(II)-5,10,15,20-tetrakis-(2,3,4,5,6-pentafluorphenyl)-porphyrin (PdTFPP) (Sigma Aldrich, USA, product number 72076-0906) dissolved in 25% w/w polystyrene (Sigma Aldrich, USA, product number 430102) in toluene was prepared and spin coated for 20 sec at 1000 RCF onto the silanized coverslips. The spin coater was maintained in a fume hood for chemical safety. The slides were allowed to dry overnight in the fume hood, protected from the light. A new set of sensor films was prepared fresh one day before each experiment.

### 3.2 Oxygen sensor film characterization

PdTFPP phosphorescence is dynamically quenched by collisions with ground state triplet oxygen.^12^ The reduction in luminescence intensity is dependent on the oxygen partial pressure according to the classic Stern–Volmer relationship (Eq 1):

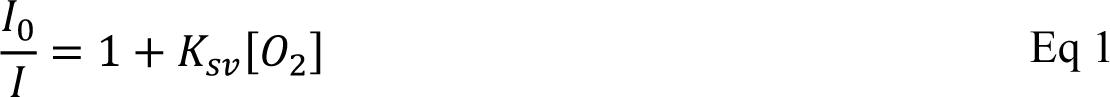

Where *I_0_* is the intensity of PdTFPP phosphorescence without a quencher at 100% N_2_ in gas phase, *I* is the intensity of phosphorescence of the sample, *K_sv_* is the quencher constant (M^-1^) and [*O_2_*] denotes the concentration of oxygen in the sample.^35^ To calibrate the oxygen sensor film, the sensor was placed in the perfusion chamber and perfused with varied mixtures of O_2_ and N_2_ in the gas phase while collecting images of the PdTFPP film (see section 3.5 Microscopy and Image processing). Images were analyzed in ImageJ. We assumed that *K_sv_* and I_0_ remained constant for all experimental measurements.

### 3.3 Lymph node slicing and staining

All animal work was approved by the Animal Care and Use Committee at the University of Virginia. Male and female C57BL/6 mice were purchased from Jackson Laboratory (USA) and used while 6-12 weeks old. Mice were housed in a vivarium and given food and water. Lymph node slices were prepared based on published protocols.^32^ Briefly, lymph nodes were embedded in 6% w/v low melting point SeaPlaque agarose (Lonza, USA) in 1× PBS without calcium or magnesium (Lonza, USA). Nodes were sliced to 300 µm thickness via Leica VT1000S vibratome (Leica, USA). Slices rested in an incubator at 37 °C for an hour before staining.

Live slices were labelled for immunofluorescence imaging according to previously published procedures.^36^ All antibodies were purchased from Biolegend. Briefly, slices were placed on a Parafilm-covered surface and a stainless-steel washer was placed on top. Samples were treated with blocking solution (anti-CD16/32) for 30 min in a cell culture incubator. Antibody cocktail (Table S2) was added to the blocking solution, and samples were incubated for an additional 1 h in the cell culture incubator. Slices were then washed by immersion in sterile 1× PBS for at least 30 min in a cell culture incubator.

For some experiments, killed control slices were generated by treatment of tissue slices with 35 % v/v ethanol for 1 hour in a cell culture incubator at 37°C, followed by rinsing with 1× PBS for 30 minutes in the incubator. Slices of inert 6 % w/v agarose were generated as controls.

### 3.4 Experimental set up for oxygen detection in tissue slices

A closed live-cell imaging chamber (Warner instruments, USA, RC-21B) was used to establish a controlled environment during imaging on the microscope stage (Figure 1). Except where otherwise noted, a gas mixture of 5 % carbon dioxide, 40% oxygen, and remainder nitrogen was bubbled into a PBS reservoir by using a pneumatic air muffler (Uxcell). The PBS reservoir was held at 37 °C in a water bath. The oxygenated PBS was perfused through the system by a peristaltic pump (Watson-Marlow, USA, 120S/DM2) with Tygon SE-200, Non-DEHP FEP-Lined tubing. An in-line solution heater (Warner instruments, SF-28) was placed immediately upstream of the imaging chamber to re-equilibrate the solution to 37 ⁰C. A flow rate of 3.6 mL/min (max speed of the pump) was used to maintain physiological temperature.

**Figure 1.**
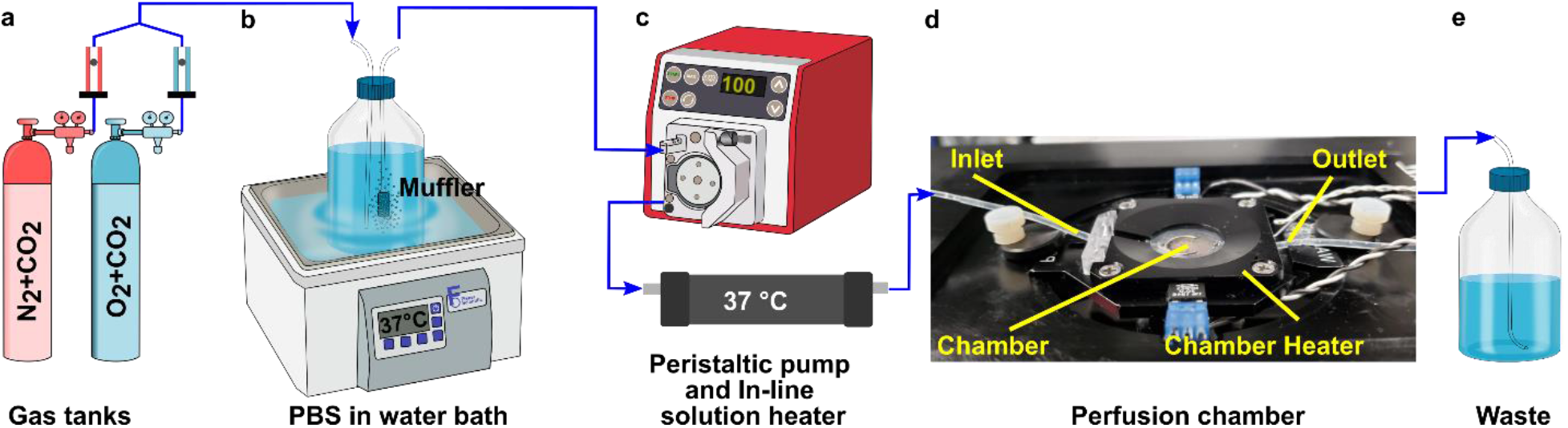
Experimental setup for imaging of oxygen consumption in tissue slices. (a) A controlled gas mixture was bubbled into (b) a PBS reservoir at 37 °C. (c) Oxygenated PBS was perfused into the closed imaging chamber using a peristaltic pump and is passed through an in-line heater (e) into the closed perfusion chamber with the sampler inside the chamber heater. (f) The excess PBS is collected in a waste jar.

To perform an experiment, lymph node slices were loaded onto a plain glass coverslip or the spin-coated oxygen sensor slide, which was quickly sealed to the imaging chamber using vacuum grease. A second plain glass coverslip was sealed to the top of the chamber to provide a closed environment. The chamber was placed in a heated stage adapter (Warner instruments, PH-2) to maintain the temperature at 37°C. Unless otherwise indicated, the chamber was perfused for 5 min after assembly with PBS bubbled with 20-40% O_2_, after which the flow was stopped to analyze oxygen consumption rate. Based on the geometry of the perfusion chamber, the shear stress applied to tissue slices at a flow rate of 3.6 mL/min was estimated at 0.146 dynes/cm^2^, which is below the reported values in lymph node tissue.^37–40^

### 3.5 Microscopy and image processing

Imaging was performed on a Zeiss AxioObserver 7 inverted fluorescence microscope with a 5X Plan-Neofluar objective, Hamamatsu ORCA-Flash4.0 LT PLUS sCMOS camera, and LED Solid-State Colibri 7 light source, which is a 7-channel LED (Zeiss Microscopy, Germany). The DAPI channel (398 nm) LED was used for excitation of the PdTFPP oxygen sensor at 50% light intensity and 150 ms exposure time, and images were collected through Zeiss penta-pass emission filter (filter set 112).

Image analysis was completed using ImageJ software 1.48v. The regional mean fluorescent intensity (MFI) of the PdTFPP oxygen sensor signal was measured in selected regions of interest, which were defined using the wizard drawing tools, in ImageJ to select the whole slice (using the brightfield images) or T cell zones (using the B220 negative regions within tissue).

#### 3.5.1 Background correction for image analysis

In most image analyses, it is sufficient to subtract a static image of the autofluorescence of the tissue. However, we found that in our timelapse imaging of the tissue on the sensor, the rate of bleaching of tissue autofluorescence and of the signal from the PdTFPP sensor were sufficient to affect the results. Therefore, we corrected the measured luminescent intensity for the effects of bleaching according to the following equation:

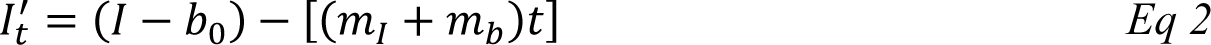

Where *I’_t_* is the corrected intensity at time *t*, *I* is the experimentally measured PdTFPP intensity, b_0_ is tissue autofluorescence at time zero, *m_b_* is the rate of photobleaching of autofluorescence of the tissue slice, and *m_I_* is the rate of bleaching of the PdTFPP sensor. The terms *m_b_* and *m_I_* were measured experimentally. The corrected intensities (*I’_t_*) were interpreted using the Stern-Volmer calibration curve (Eq. 1) and converted from mmHg to mM of dissolved O2 by using Henry’s law with a Henry’s law constant of 767.69 mm Hg/mM (see SI Methods).

### 3.6 Predicting the uncertainty in oxygen concentration measurements

The measured intensity values (*I*) at any given location are the sum of tissue autofluorescence and PdTFPP signal at that location. Uncertainties in these values were measured experimentally (see section 4.4) at the start of the measurement (*t* = 0 s). As these were independent sources of variation, error in the background-corrected *I’_t=0_* values were calculated as follows:

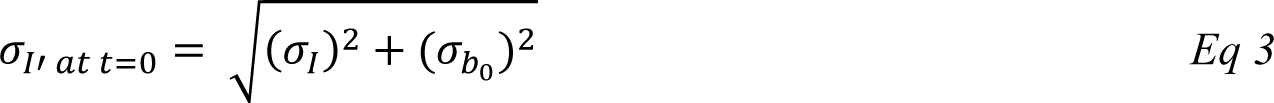

Where *σ_I_* is the standard deviation of the intensity of the oxygen sensor film at different locations, and *σ_bo_* is the standard deviation of repeated measurements of tissue autofluorescence intensity, both prior to photobleaching.

To estimate the error in the measurements of [*O_2_*] calculated from the Stern-Volmer equation (Eq 4), we used the general error propagation formula (Eq 5) to calculate the uncertainty in [*O_2_*] measurements (Eq 6). Here, *σ_I’_* _at t=0_ was determined from Eq. 3, and *K_SV_* and *I_0_* were treated as constants.

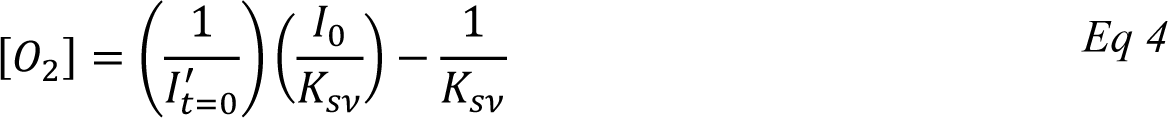

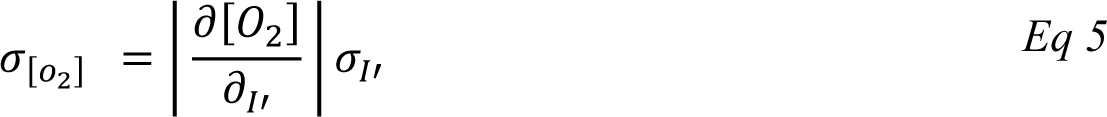

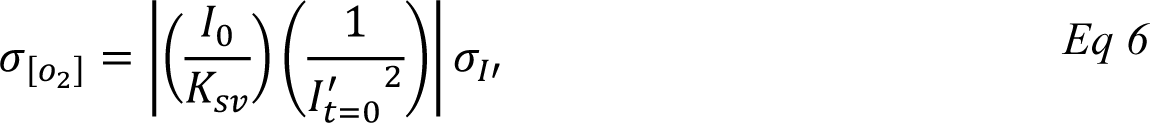

### 3.7 Seahorse metabolic analysis

T cells and B cells were isolated from murine lymph nodes using EasySep™ Mouse T cell Isolation Kits or B cell Isolation Kits (negative selection; STEMCELL Technologies, USA) according to manufacturer instructions. Metabolic analysis was conducted on a Seahorse Xfe96 analyzer (Agilent) according to published procedures.^41^ Briefly, cells were plated at 300,000 cells/well in XF96 plate. Keeping the temperature constant at 37°C by pre-warming all media was especially important to a successful experiment. OCR was measured using the mitochondrial stress test procedure, using XF media that consisted of non-buffered DMEM (Agilent, USA) supplemented with 10 mM glucose (Gibco, USA), 2 mM L-glutamine (Gibco, USA), and 1 mM sodium pyruvate (Gibco,USA). The Mito-stress test was performed according to manufacturer instructions in response to 1 µΜ oligomycin (Sigma Aldrich), 1 µΜ fluoro-carbonyl cyanide phenylhydrazone (FCCP) (Sigma Aldrich), and a mixture of 0.5 µΜ rotenone (MP Biomedicals), and antimycin (Sigma Aldrich).

## 4 RESULTS

### 4.1 Analytical set up and sensor characterization

To visualize and quantify the rate of oxygen consumption in live tissue slices ex vivo, we designed a system to collect timelapse images of an O_2_-responsive glass sensor under a tissue sample under well controlled culture conditions (Figure 2). We selected an oxygen sensing film previously described by the Lockett laboratory at University of North Carolina, which is generated by mixing palladium tetrakis(pentafluorophenyl)porphyrin (PdTFPP) into a polystyrene matrix.^24^ Porphyrin dyes, known for their high brightness and sensitivity to oxygen, are commonly used in oxygen sensing studies.^42^ PdTFPP in particular offers a wide range of oxygen detection from 0 to 100% oxygen and is compatible with standard DAPI excitation and emission filters,^43^ and PdTFPP films exhibit a linear and highly sensitive response to changes in oxygen tension.^24^ Here, we prepared an oxygen sensor by spin-coating PdTFPP dye in polystyrene onto a glass coverslip.

**Figure 2.**
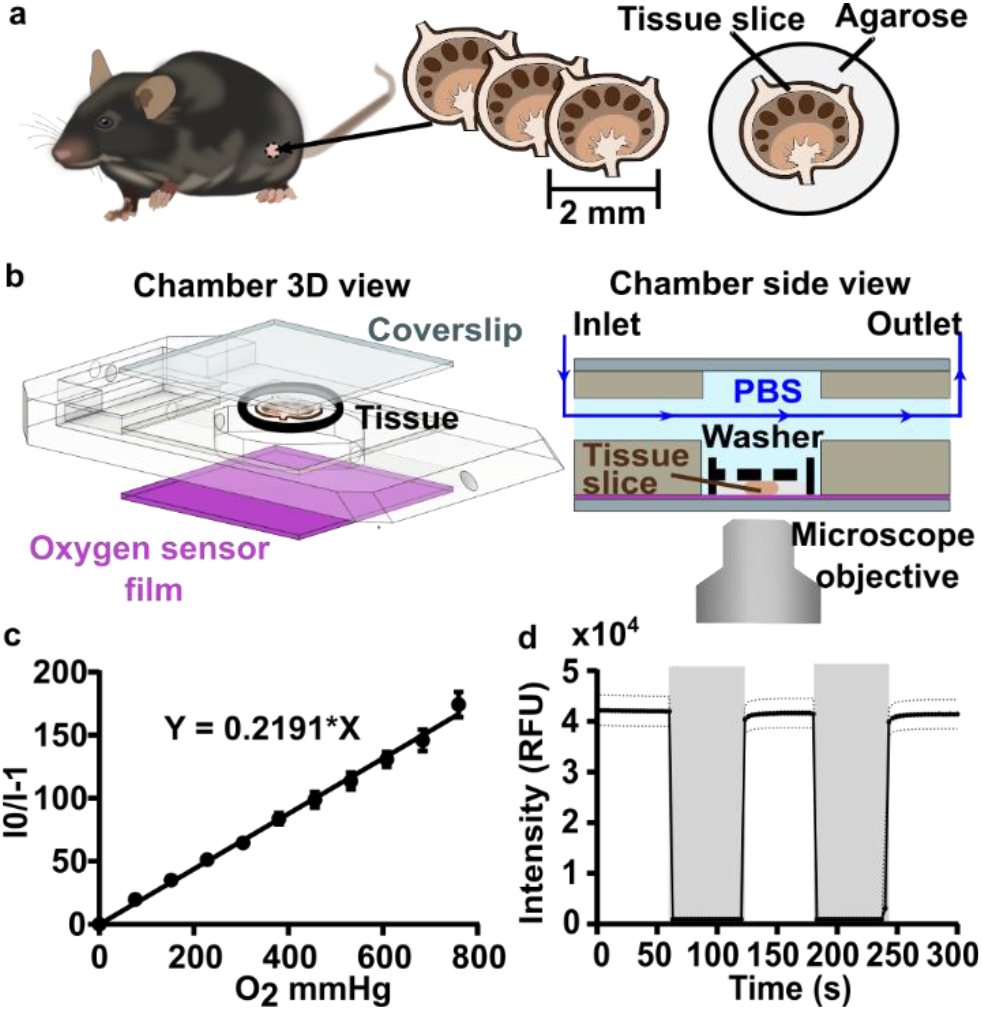
Analytical system set up and sensor characterization. (a) Murine lymph nodes were harvested and sliced using a vibratome. (b) CAD drawing (exploded view, left) and schematic (side view, right) of the perfusion chamber and assembled imaging setup, respectively. Oxygen-bubbled PBS or gaseous media were flowed through the chamber. The slice was imaged with or without flow as indicated during each experiment. A small washer (black dashed line) kept the lymph node slice in contact with the bottom coverslip (purple, sensor film). (c) Gas-phase calibration curve as a function of oxygen partial pressure. A linear fit according to Stern-Volmer theory yielded a K_sv_ of 0.219 ± 0.001 mmHg^-1^. Each data point is the average of 3 measurements from three independent films; error bars represent standard deviation. (d) Response characteristic of the PdTFPP oxygen sensor film in a gas-filled chamber that was flushed alternately with 100% oxygen (grey shading), and 100% nitrogen (no shading). Each data point is an average of 3 measurements from three independent films; dotted lines represent standard deviation.

Next, we selected a closed-bath perfusion chamber based on its ability to incorporate the sensor-coated glass from the bottom, seal the chamber from the outside air to control the interior gas content, and accommodate the size of the tissue slice (300 µm thickness and 10 mm diameter) with head space for gas or buffer flow above. The sample was loaded onto the sensor slide and the chamber quickly assembled, connected to gas or buffer flow, and imaged on the microscope (Figure 2b). This system provided control over the tissue environment during imaging, including oxygen content, temperature, and flow rate.

We first confirmed that the oxygen sensing film responded as expected to oxygen partial pressure in the gas phase, in the absence of tissue. As expected, the intensity of the sensor was inversely correlated with partial pressure of oxygen in the gas phase, and the Stern-Volmer plot was linear with a K_sv_ value of 0.2191 ± 0.001 mmHg O_2_ ^−1^ (Figure 2c), similar to reported values.^24^ The signal from PdTFPP exhibited a reasonable level of photostability, showing a 0.8% decay during a ten minute imaging period in the gas phase (SI Figure 1). As we were interested in the dynamics of oxygen consumption, we characterized the response time and stability of the sensor under oscillating oxygen partial pressure of 0 and 100 % oxygen in the gas phase. The intensity returned to the same value with each oscillation, indicating stability and lack of hysteresis. Furthermore, the response time of PdTFPP to changing oxygen environment was less than 3 seconds (Figure 2d), which was considerably faster than the time needed for consumption of oxygen in subsequent tissue experiments.

### 4.2 Spatially resolved oxygen mapping revealed greater depletion of oxygen in T cell zone than elsewhere in lymph node tissue

As a first test of oxygen consumption in live, metabolically active tissue, we compared the response of the sensor to live lymph node slices versus ethanol-treated tissue and inert slices of agarose gel. After loading the sample into the chamber, we stopped the perfusion of PBS to isolate the sample from any replenishment of oxygen, and then imaged the oxygen levels that remained after a short incubation time (5 min). The PTFPP signal became bright in the chamber housing live tissue, indicating depletion of oxygen, and remained negligible in chambers housing the negative controls, as expected (Figure 3a). By averaging the intensity across the tissue and using the Stern-Volmer calibration to quantify mean oxygen concentration, we found an equivalent of 172 ± 5, 164 ± 28, and 10 ± 3 µM O_2_ remaining in the agarose, killed and live slices, respectively (Figure 3b). The former values are roughly consistent with the expected dissolved oxygen level in media incubated at 37°C, which is 0.181 mM at sea level.^44^

**Figure 3.**
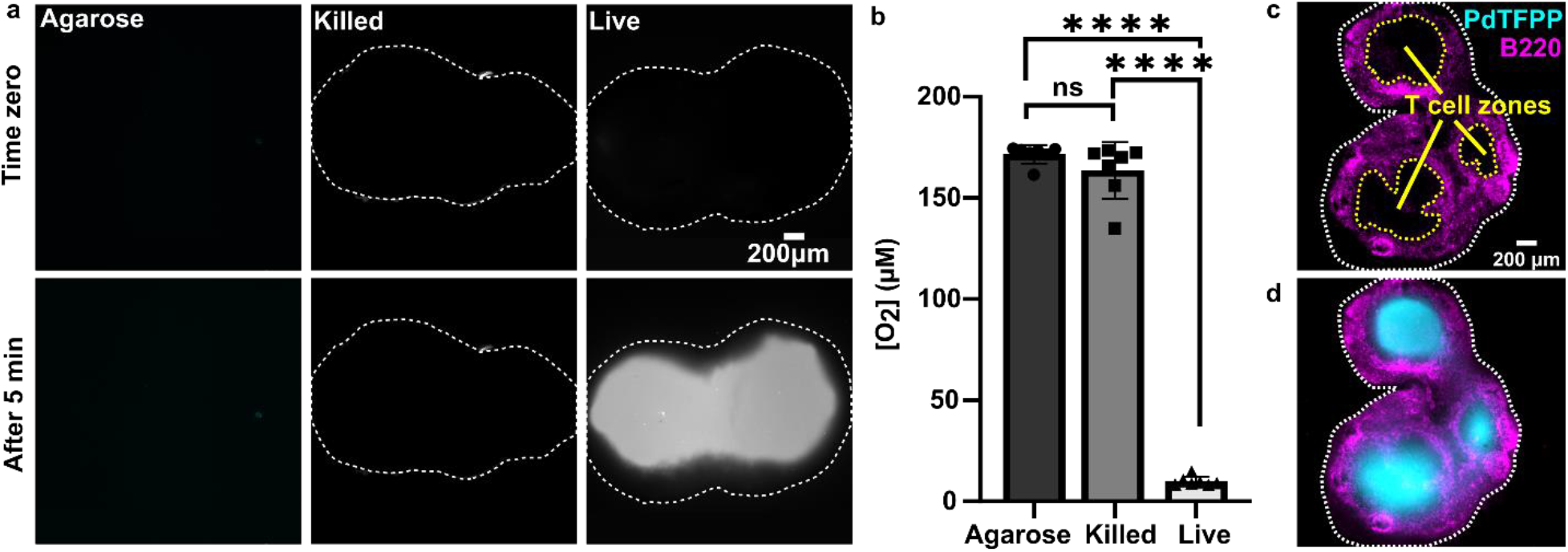
Detection of local oxygen depletion by live lymph node slices. (a) Greyscale images of PdTFPP signal from agarose slices, killed tissue, and live tissue that were loaded into the chamber, perfused briefly with PBS, imaged immediately after the flow of PBS was stopped (time zero), and imaged again five minutes later. Oxygen was consumed in the live slice after 5 minutes (high PdTFPP signal) while there was no consumption in killed or agarose slices. The white dotted lines denote the outline of the tissue slice. The PBS was not bubbled with oxygen in this experiment. (b) Graph of mean oxygen concentration in the sample, quantified from images collected after 5 minutes. Error bars denote standard deviation. N = 6 slices per condition, one-way ANOVA analysis with Tukey post-hoc test, **** indicates p < 0.0001. (c,d) Representative images of a lymph node slice immuno-stained for B220 (B cell marker, fuchsia) (c) The yellow dashed lines indicate the borders of the T cell zone regions of interest that were defined for subsequent analysis (see SI). (d) with the signal from the PdTFPP sensor (cyan) overlaid.

Interestingly, in the live samples, we consistently observed that the periphery of the tissue had much dimmer PdTFPP signal than the center of the tissue (Figure 3a), suggesting less oxygen consumption in the peripheral regions. The lymph node is a highly organized organ, with B cell follicles and lymphatic sinuses located near the exterior, and T cells enriched in a central paracortical region.^45^ To map this distribution of signal against the regional tissue organization, we combined the oxygen sensing method with live immunofluorescence labelling of slices for the B cell marker B220 (Figure 3c).^36^ In addition to B cell follicles, this antibody also labels the sinuses at a lower intensity through Fc-mediated binding, thus highlighting the outer cortex and leaving the interior T cell zone dark.^36^ By labeling the lymph node slices before placement into the chamber for the oxygen depletion assay, we found that the PdTFPP signal was consistently localized to the central, B220-negative regions (Figure 3c). Thus, oxygen was consumed far more rapidly in the T cell region than in the B cell region of the lymph node tissue. This pattern was remarkably consistent across heterogeneous tissue samples (SI-Figure S2).

### 4.3 Temporal analysis of rate of oxygen consumption in the T cell zone of lymph node slices

During the above experiments, oxygen consumption occurred within 1 to 5 minutes upon assembly of the perfusion chamber, and was often partially or fully complete before imaging began. To enable quantification of the initial rate of consumption, we attempted to replenish the oxygen that was consumed during assembly by perfusing the chamber with oxygen-bubbled PBS for 5 minutes prior to stopping the flow and imaging the PdTFPP over time (Figure 4a). Using this protocol, we were able to observe oxygen consumption by lymph node slices in real time in most tissue slices (Figure 4b and Supporting movies).

**Figure 4.**
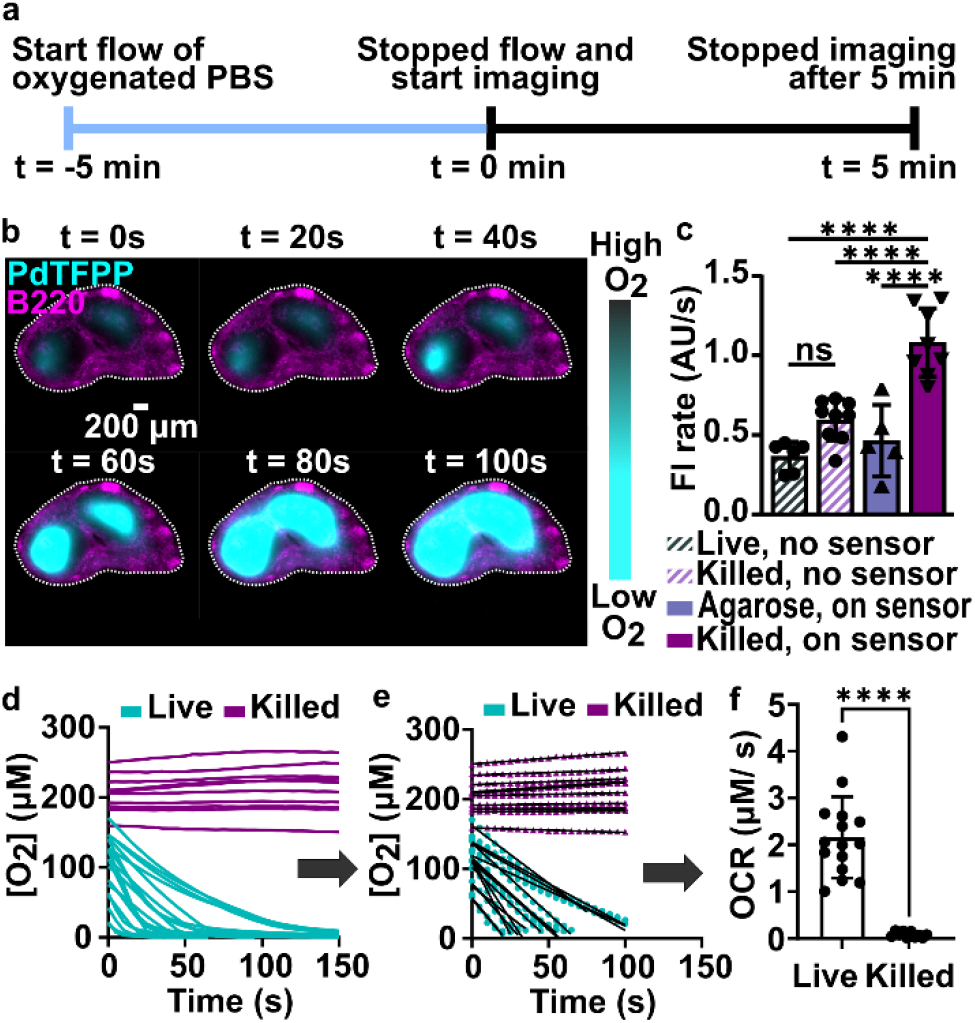
Spatially resolved quantification of oxygen consumption rates in lymph node slices. (a) Timeline for sample loading and imaging. The stained slices were loaded onto the chamber, and the chamber was connected to oxygen bubbled PBS (t = -5 min). The flow was stopped at t = 0 min, and slices were imaged every 5 seconds for 5 minutes. (b) Timeseries of fluorescence images of a representative lymph node slice immuno-stained with B220-Alexa Fluor 647 for B cell follicles (magenta) overlaid with the signal from the PdTFPP oxygen sensor (cyan). White dotted lines indicate the outline of the tissue slice. Brightness and contrast were uniformly adjusted between images. (c) Plot of rates of photobleaching of the PdTFPP oxygen sensor and autofluorescence of tissue. N = 6 samples per condition, one way ANOVA analysis with Tukey post-hoc test, ns indicates no significance and **** indicates p < 0.05. (d) Representative plot of oxygen concentration over time in regions of interest in live and killed tissue slices. Each line indicates a distinct T cell zone in total of 12 slices per condition, pooled from 1 male and 1 female mouse. (e) The initial, linear portion of the data in (d) was fit to determine the initial rate of oxygen consumption. (f) Comparison plot of the initial rate of oxygen consumption in live and killed slices calculated from the slope of the linear fit as demonstrated in (e), pooled from repeated experiments. Each dot indicates a distinct T cell zone in total of 12 slices per condition n of 2 mice. Student T test with unpaired two tailed analysis, **** indicates p < 0.0001. Error bars denote standard deviation.

In initial experiments using this protocol, we observed that both the autofluorescence of the tissue and the signal from the PdTFPP sensor were bleaching slowly during the imaging time course, leading to falsely low fluorescent signals and thus falsely high calculated oxygen concentrations at later times. To correct for the effect of these two factors, we designed a pair of experiments to quantify the individual photobleaching rates in the absence of oxygen consumption (Figure 4c). To quantify the rate of photobleaching of lymph node tissue autofluorescence, we used live slices or ethanol-treated slices, which are not metabolically active, and quantified the change in tissue autofluorescence in the PdTFPP channel without the PdTFPP sensor during repeated imaging. The two rates were not significantly different, so we made a simplifying assumption that all live and killed tissue samples would bleach at the same constant rate (0.7 ± 0.09 AU/s). Next, to quantify the bleaching rate of PdTFPP, we measured bleaching of the PdTFPP film by loading agarose slices into the chamber instead of tissue and again quantified intensity over time. The rate of PdTFPP oxygen sensor bleaching was 0.4 ± 0.2 AU/s. Both the tissue autofluorescence and PdTFPP sensor photobleached linearly over time. After confirming that these rates combined additively by quantifying the bleaching of killed slices on the sensor, we used them to correct the observed fluorescent intensity (Eq 2) prior to calculating [O_2_] from the Stern-Volmer calibration curve. This correction yielded essentially stable [O_2_] in the chamber with killed slices over time (Figure 4d), as expected, and was used for analysis of all tissue slices for the remainder of the paper.

Since most consumption occurred in the T cell region, we specifically analyzed the oxygen consumption rate (OCR) in individual T cell regions, rather than averaging across the whole slice. We fit the initial portion of each data set (Figure 4d) with a linear regression (Figure 4e), yielding an OCR in T cell zones of live and killed slices of 2.16 ± 0.87 and 0.08 ± 0.06 µM/s in T cell zones of live and killed slices, respectively (mean ± stdev, n = 12). As a practical matter, it was not possible to obtain reliable initial rates in T cell regions that had initial oxygen concentration of < 0.05 mM at t = 0 (when flow was stopped), so these samples were excluded from analysis of initial rates. This limitation potentially excludes the fastest consuming samples from the data set, amounting to approximately 1 or 2 of slices out of n = 12. Nevertheless, this method enabled spatially quantification of oxygen consumption rates in the majority of samples, which has not been possible previously.

Finally, we note that although oxygen must be consumed by cells in the B cell region, in these naïve lymph node slices its rate was sufficiently slower than that of the T cell zone that it was not quantifiable by this method. The region of depleted oxygen spread by diffusion from the rapidly consuming T cell zone into the B cell area within minutes (in other words, oxygen diffused from the B cell zone into the T cell zone). In future studies, modifications to the method would be needed to quantify oxygen consumption rates in slow regions that are adjacent to rapidly consuming regions. Interestingly, B cells experience a dramatic increase in basal and maximal OCR upon activation, even exceeding that of activated T cells,^46^ suggesting that it may be possible to detect oxygen consumption by activated B cells in reactive lymph node tissues.

### 4.4 Variation in initial oxygen concentrations is largely due to measurement uncertainty

Unexpectedly, we observed substantial variability in initial oxygen concentration (time zero in Figure 4d), ranging from 0.06 to 0.19 mM for live slices, with a standard deviation of 0.03 mM O_2_ (data from n = 2 mice and 12 slices). We reasoned that this variability could be due to either (i) random variations in the measurement, particularly because high initial [O_2_] corresponds to a dim signal from the PdTFPP dye (low signal/noise), or (ii) biologically meaningful variations in the rate of oxygen consumption prior to starting perfusion, such that slices that consumed very rapidly would never “recover”.

To isolate the contribution of random measurement variation, we used ethanol-treated slices that could not consume oxygen, and used error propagation to determine the uncertainty in the initial oxygen concentration, [O_2_]_t=0_ (Section 3.6 in Methods). [O_2_]_t=0_ is calculated from the Stern-Volmer equation (Eq 4) using the autofluorescence-corrected intensity *I’* at time zero, which depends solely on the PdTFPP signal (*I*) and the tissue autofluorescence (*b_0_*) (Eq 2). Uncertainty in *b_o_* (σ*_bo_*) was determined from the standard deviation of the intensity of autofluorescence measured at multiple locations in killed tissue samples under perfusion of oxygenated PBS, in the absence of the PdTFPP sensor (three locations per slice and N = 6 slices). Uncertainty in measurements of *I* (*σ_I_*) was determined from the standard deviation of the intensity of the PdTFPP sensor measured at multiple locations of agarose slices under perfusion of oxygenated PBS (three locations per slice and n = 6 slices; n = 3 films per sample). Propagation of error (Eq 6) yielded a predicted uncertainty of 0.02 mM O_2_, which was of the same order of magnitude as the measured variation in [O_2_]_t=0_ in live slices (0.03 mM). Furthermore, there was no correlation between [O_2_]_t=0_ and the initial rate of consumption (SI Figure S3), suggesting that a lower [O_2_]_t=0_ was not attributable to faster consumption prior to the start of the measurement. We thus concluded that random variation was sufficient to explain at least the majority of the observed variation in initial oxygen concentration, and that this variability did not affect the quantification of rate of consumption in slices above the sensitivity threshold.

### 4.5 Validation of regional oxygen consumption rates compared to Seahorse instrument

We sought to test the accuracy of the optical method by comparing its results to those of the gold-standard Seahorse method (XFe96 flux analyzer). The Seahorse assay measures the oxygen concentration in cell culture supernatants over time to determine OCR (pmol O_2_/min/cell). As it was not possible to load the 300-µm thick lymph node slices into the Seahorse instrument, we compared against cell suspensions isolated from naïve murine lymph nodes. First, building on our observation that oxygen consumption was significantly faster in the T cell region than in the B cell region, we measured the OCR for purified T cells and B cells by using a mitochondria stress test assay on XFe96 Seahorse instrument (SI Figure S4). Consistent with prior reports,^47^ T cells had a significantly higher basal OCR than B cells (Figure 5a,b); this result may partially explain the spatial distribution of oxygen consumption in the naïve lymph node slices in the optical assay. Next, to validate quantitative accuracy, we compared the OCR measured by the two methods. To convert the rates of oxygen consumption per T cell zone from the optical method (mM/s) to an order-of-magnitude estimate of OCR per cell (pmol/min/cell), we assumed an average number of T cells per slice (SI assumptions and calculations), an average of two T cell regions each of average area, and fixed tissue thickness (300 µm).^32^ These simple estimates yielded an average basal OCR that was within 2-fold and not significantly different from that obtained from purified T cells using the Seahorse instrument (Figure 5b and SI calculations). The consistency between the tissue-based and cell suspension-based assays suggests that in the lymph node slice, oxygen consumption of lymphocytes likely exceeds that of stromal and antigen presenting cells, as observed previously for glucose consumption,^33^ although we did not test that here. In summary, the optical method in tissue provided OCR similar to standard in vitro cell culture measurements, while also providing regional spatial resolution in tissue slices.

**Figure 5.**
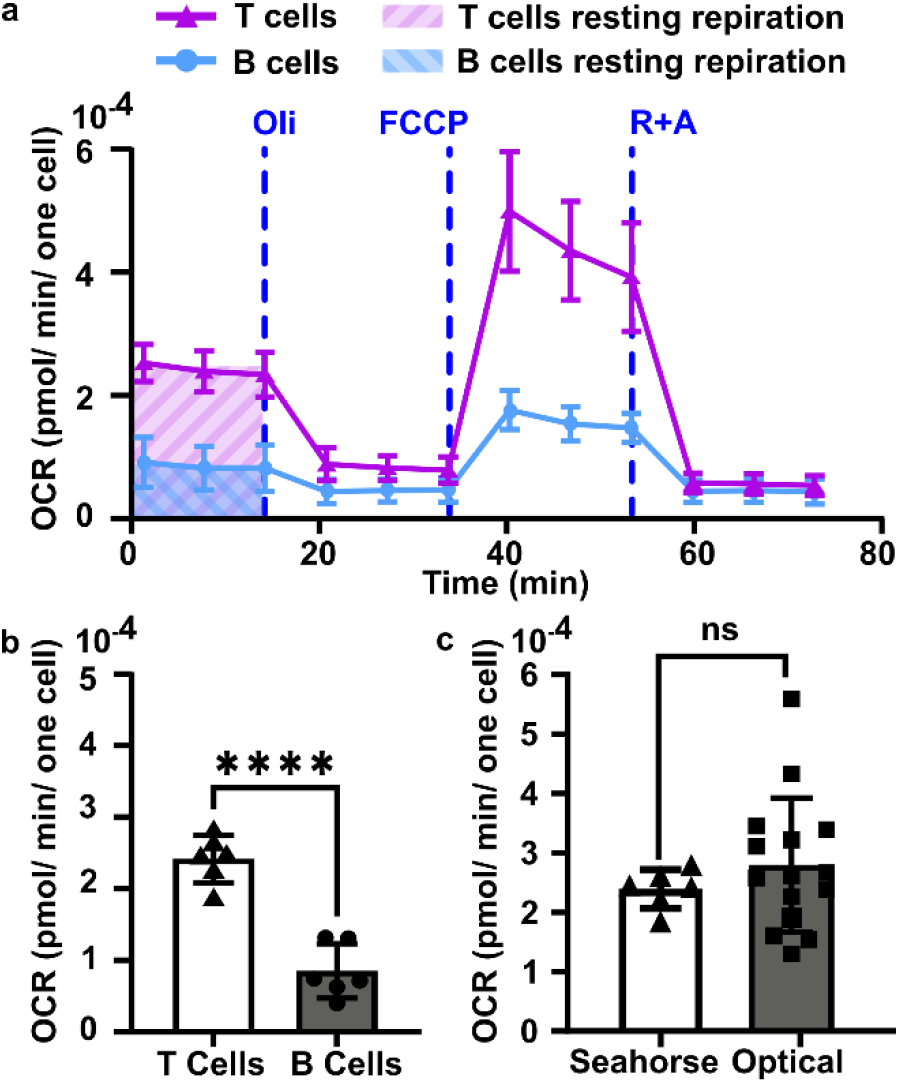
Quantification of oxygen consumption in lymph node slices compared to Seahorse method with cell suspensions. (a) Representative metabolic flux analysis of B cells and T cells isolated from murine lymph nodes. Cells were analyzed in microplates by Seahorse XP 96 according to the Mito Stress protocol with 300,000 cells per well. Where blue dashed lines are indicated, the following compounds were injected into the assay micro-chambers: oligomycin (Oli), carbonyl cyanide-4-(trifluoromethoxy)phenylhydrazone (FCCP), rotenone plus antimycin A (R+A). (b) Average basal rate of oxygen consumption of T cells and B cells from naive murine lymph nodes, measured by Seahorse Mito-stress test. Student T test with unpaired two tailed analysis, **** indicates P < 0.0001, N = 6 wells (pooled from 1 male and 1 female mouse). (c) Comparison of rate of oxygen consumption measured by Seahorse and optical methods, using isolated T cells for Seahorse and the T cell zone region of interest in lymph node slices for the optical method from figure 4 (Each dot is one T cell zone, from N = 12 slices for optical method; N = 6 wells for Seahorse, pooled from one female and one male mouse. All cells and tissues were collected from naïve animals. Student T test with unpaired two tailed analysis, ns indicates not significant (p > 0.05). All bars denote mean and standard deviation.

### 4.6 Continuous perfusion replenishes oxygen to prevent lymph node hypoxia

Tissue slices are often cultured either under continuous perfusion, on a rocking platform to induce fluid mixing, or at an air-liquid interface. Each of these strategies is intended to improve oxygenation of thick tissue samples, but rarely is the effect of the intervention on O_2_ levels directly measured. Here, as a proof of principle of the utility of the assay, the optical oxygen sensing assay was employed to directly measure the impact of continuous perfusion on oxygen depletion by lymph node slices in a closed chamber. After assembling the chamber and perfusing for 5 min to equilibrate as above, each slice was imaged an additional 5 min either under continuous flow of oxygenated PBS or under static conditions. To increase the power of the experiment, each tissue was tested twice, once for each condition.^33^ As expected, the slices became significantly more hypoxic under static conditions than under continuous perfusion (Figure 6). The mean rate of oxygen consumption in the T cell zone without flow was 1.6 ± 1.0 µM/s, versus just 0.30 ± 0.28 µM/s for slices with flow (N = 12 for each group; Figure 6b). Continuous perfusion of oxygenated PBS maintained stable oxygen levels, indicating an adequate oxygen supply with this flow rate (3.6 mL/min). Thus, the optical assay was able to easily distinguish between the two conditions, highlighting its utility to guide the selection of culture conditions for cell and tissue culture.

**Figure 6.**
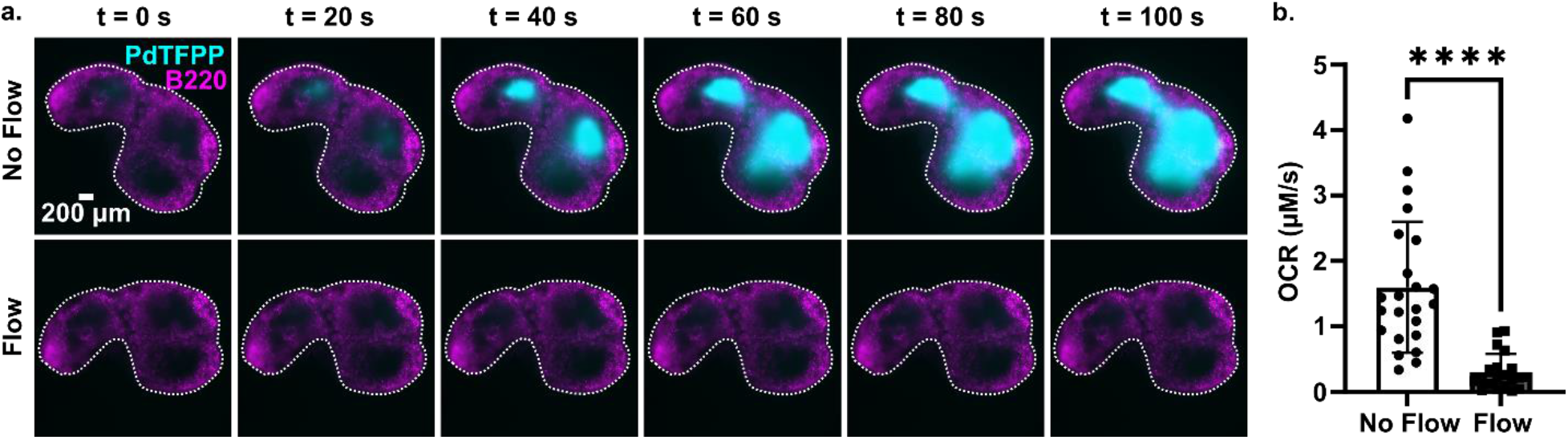
Application of the assay to quantify the effect of fluid perfusion on prevention of hypoxia in ex vivo lymph node slices. (a) Timelapse images of PdTFPP signal from slices cultured under static conditions or under continuous perfusion. Lymph node slices were immuno-stained with B220 Alexa Fluor 647 for B cell follicles (magenta); PdTFPP oxygen sensor shown in cyan. Bright PdTFPP signal indicates low local oxygen concentration. White dotted lines indicate the outline of the tissue slice. Brightness and contrast were uniformly adjusted between images. (b) Plot of initial oxygen consumption rate in slices under static or continuous perfusion. N = 12 lymph node tissue slices per condition. Each dot indicates one T cell zone. Student T test with paired two tailed analysis **** indicates P < 0.0001. Error bars denote standard deviation.

## 4 CONCLUSIONS

In summary, we report a quantitative imaging method that allows mapping of the spatial distribution of oxygen consumption rate in ex vivo tissue slices. In contrast to measurement of OCR in cell suspensions, the ex vivo approach maintains the tissue microenvironment, allowing for the examination of region-specific cellular metabolic activity. Correction for photobleaching of both the tissue and the oxygen-sensitive dye was required for accurate quantitation of consumption rates. This method correlated well with the standard Seahorse method, while also revealing for the first time that in naïve lymph node slices, the T cell zone consumed oxygen far faster than the cortical and B cell region. Repeated testing of the same tissues was possible, thus increasing the power of experiments, and we used this approach to evaluate the impact of continuous on prevention of hypoxia in individual tissue slices. In addition to testing the impact of varied culture conditions, we anticipate that this method will be useful to analyze regional oxygen consumption in diseased tissues and to assess the impact of drug therapy. Furthermore, there is potential to integrate this method with other existing assays, such as glucose uptake assays,^33^ to gain a more comprehensive understanding of the metabolic activity of in live tissues.

## ACKNOWLEDGEMENTS

This work was supported by the National Institute of Allergy and Infectious Diseases (NIAID) of the National Institutes of Health (NIH, award numbers R01AI131723 and 1R21AI160547). The content is solely the responsibility of the authors and does not necessarily represent the official views of the National Institutes of Health. We would also like to thank Cassandra Fraser (Univ. of Virginia) and Mathew Lockett (Univ. of North Carolina, Chapel Hill) for their insight and guidance for selecting the oxygen sensor. The authors thank Drake D. Dixon (Univ. of Virginia) for experimental contributions at the early stages of this project.

