## Supplemental information for "Spatially resolved quantification of oxygen consumption rate in ex vivo lymph node slices"

**Supplemental Figures**

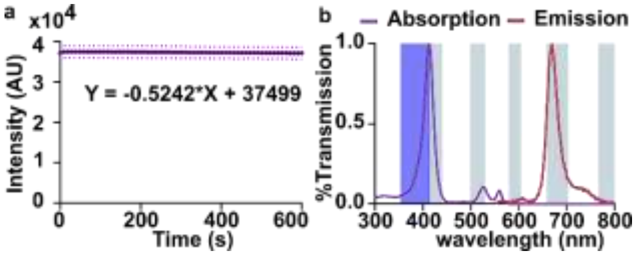

**Figure S1: Characterization of the photostability and absorption and emission spectra of the PdTFPP sensor.** (a) Luminescent intensity (arbitrary units) of the PdTFPP oxygen sensor film in gas phase 100% nitrogen over 45 min. Images were collected every 10 s at 150 ms exposure time; the shutter was closed between exposures. (b) Absorption (purple) and emission (red) spectra of PdTFPP. The wavelengths of the pentapass emission filter (grey; Zeiss Filter set LED 112) and of the DAPI excitation LED (dark blue) are also shown.

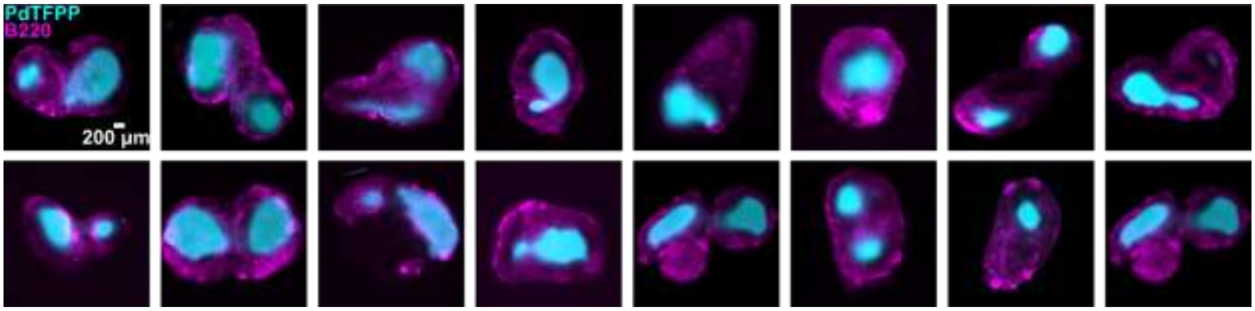

**Figure S2: Distribution of oxygen consumption in lymph node slices from naïve mice.** Each image was collected 5 minutes after the lymph node tissue slice was placed onto the perfusion chamber and the flow of oxygenated PBS was stopped. Oxygen was consumed mostly in T cell regions of the slices (PdTFPP oxygen sensor in cyan). The B cell regions were stained with B220 (magenta).

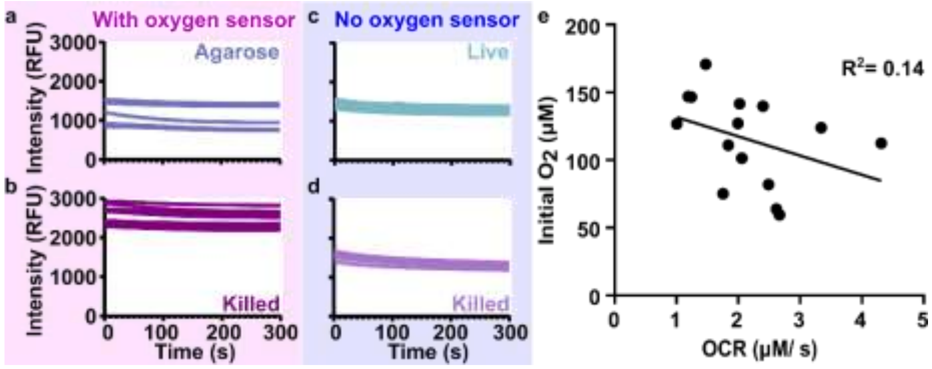

**Figure S3: The photobleaching rates of PdTFPP oxygen sensor film and of tissue autofluorescence in oxygenated PBS.** (a,b) Plot of fluorescence signal from the PdTFPP oxygen sensor over time during imaging of (a) an agarose slice or (b) an ethanol-killed tissue slice. The shutter was closed between the takes. (c,d) Plots of fluorescence signal from (c) live and (d) ethanol-killed slices without the oxygen sensor over time. N = 6 slices for all conditions; each curve shows one slice. Same y-axis scale as in (a,b). (e) Correlation plot between the initial $O_2$ concentration and the initial rate of $O_2$ consumption in live tissue slices, for the data shown in Figure 4e. The strength of a linear association between the two variables was measured using a two-tailed Pearson correlation coefficient test, and the R squared value of 0.14 indicated no correlation between the variables.

**Captions for supplemental movies**

Movies showing fluorescent signal from PDTFPP oxygen sensor overlaid with live and killed lymph node tissue slices. Images were taken every 5 seconds for a total of 5 minutes. PdTFPP signal is in cyan color and B220 (B cell biomarker) is in magenta color. Three movies are provided; two representative of live slices and one for killed slices.

### Supplemental Tables

**Table S1: Fluorescent antibodies used to label cells for fluorescence microscopy.**

| Biomarker/ reagents | Clone | LOT | CAT | Vendor |
|---|---|---|---|---|
| CD16/32 | 93 | B298973 | 101302 | Biolegend |
| B220 | RA3-6B2 | B243962 | 103226 | Biolegend |

### Supplemental assumptions and calculations

*Dissolved oxygen calculations and assumptions: Converting mmHg to mM $O_2$*

Dissolved oxygen concentration in solution is affected by factors such as atmospheric pressure and the solubility of $O_{2(g)}$ in the liquid, which is inversely proportional to solvent temperature and ionic strength. To calculate the solubility of oxygen gas in PBS, we used Henry's law of solubility (Eq. S 1). This Law provides a mathematical description of gas solubility in a liquid medium. According to Henry's law, at a constant temperature, the solubility of a gas in a liquid is directly proportional to the partial pressure of the gas above the liquid at equilibrium.

$$C = \partial P_{gas} / H \qquad \text{Eq. S 1}$$

Where $C$ is the solubility of a gas at a fixed temperature in a particular solvent (mM), H is Henry's law constant (mmHg/ mM), and $\partial P_{gas}$ is the partial pressure of the gas (mmHg). The Henry's law constant for a solution is dependent on the concentration of electrolytes and proteins, atmospheric pressure, and temperature of the solution. Here, we assumed a standard atmospheric pressure of 760 mmHg and temperature of 37 °C for all experiments. Ionic strength of the 1x PBS used here (LONZA; Catalog No: 17-516F) was calculated to be 166 mM at pH 7.4.

Next, we determined the partial pressure of $O_2$ in the cell culture incubator and in the stage-top chamber used for the optical assay. At 37 °C and 100% humidity (inside incubator), water vapor exerts a partial pressure of 47 mmHg.[1] Therefore, water vapor will make up 6.2% of the total gas at sea level (47 mmHg/ 760 mmHg). To calculate the final $O_2$% in the gas phase under different conditions used for cell culture, we used the following equation:

$$\%O_2\,final = (\%O_2\,initial \times (1 - \%gas_{(1)} - \%gas_{(n)} + \dots)) \times 100 \qquad \text{Eq. S 2}$$

Therefore, for the following conditions we have the following sample calculation:

- Humidified incubator without 5% CO2 (Place et al. 2017)

  $\%O_2$ final = [%O2 initial$_{(atmospheric)}$ × (1- % $H_2O$ vapor)] × 100

  $\%O_2$ final = [0.21 × (1 – 0.062)] × 100 = 19.7%

  $PO_2$ = 760 mmHg × $\%O_2$

  $PO_2$ = 760 mmHg × 0.197 = 149.7 mmHg

The extrapolated dissolved oxygen concentration at ionic strength of 166 mM in a humidified incubator without $CO_2$ is 0.195 mM (Place et al. 2017, Supplemental Table 1).[2–4] Henry's constant under these known conditions can be solved as follows:

- Incubator without 5% $CO_2$

  $H = \partial P_{gas}/C$ = 149.7 mmHg / 0.195 mM; H = 767.69 mmHg/mM

Therefore, we used 767.69 mmHg/mM as the Henry's constant for conversions between mmHg to mM $O_2$ throughout the manuscript.

*Shear stress calculation for flow of PBS in perfusion chamber*

To estimate the fluid shear stress (*FSS*) resulting from the flow of 1x PBS through the perfusion chamber, we assumed a thin rectangular cross section (Eq. S 3)[5]:

$$FSS = (6\eta Q) / (h^2 \times w) \qquad \text{Eq. S 3}$$

Where η is fluid viscosity, $Q$ is fluid flow rate, $h$ is height of the bath, and $w$ is width of the bath. For the closed bath chamber, we have $Q$ = 3.6 mL/min = 0.06 $cm^3$/s, $\eta$ of PBS = approximately 1.00 × $10^{-3}$ Pa·s,[6] $h$ = 0.25 cm, and $w$ = 1.3 cm (Warner Instruments). Thus, the fluid shear stress is calculated as

$$FSS = \frac{[(6)(1.00 \times 10^{-3}\ Pa.s)\left(0.06\frac{cm^3}{s}\right)]}{(0.25\ cm)^2\ (1.3\ cm)} = 0.004\ Pa$$

Converting from Pa to dyn/$cm^2$ (1 Pa = 10 dyn/$cm^2$), we have an estimated *FSS* of 0.044 dyn/$cm^2$ in the chamber under these conditions.

*Rates of oxygen consumption: Assumptions and calculations*

At each time point, the fluorescence intensities from the oxygen sensor film were collected in mean grey value, which was converted to [dissolved $O_2$] (mM) by using the oxygen calibration curve for the perfusion chamber (Stern-Volmer equation, Eq. 1 from main text) and Henry's law. After generating a plot of $[O_2]$ (mM) vs time (s), the initial portion of the curve was fit with a linear fit, and the mean rate of consumption per tissue slice (mM/s) was obtained from the absolute value of the slope (Figure 4d-f).

To compare the optical assay to the results from a Seahorse assay, it was necessary to convert between units of mM/s per T cell zone and pmol/min/cell. Doing so required a series of assumptions and simplifications, yielding an order-of-magnitude comparison between the two measurements. To convert from molarity to moles by using the molarity equation (Eq. S 4),

$$\textit{Molarity} = \textit{moles of substrate} / \textit{volume of solution} \qquad \text{Eq. S 4}$$

We estimated the average volume (area x height) of the T cell zone in a lymph node slice. To determine an average area, we performed immunofluorescence labelling of lymph node slices, defined the T cell zones as central B220-negative regions (see e.g. Figure 3c from main text), and measured the areas of these regions using ImageJ. Measurements in 12 naïve lymph node slices from 6 wks old male and female mice yield an average area of $1.01\times 10^6 \pm 0.26 \times 10^6$ μm$^2$. Assuming a thickness of 300 μm, this yielded an average volume of $3.02 \times 10^8$ μm$^3$.

To convert from moles per T cell zone to moles per cell, we estimated an average number of T cells per T cell zone. To do so, we referred to the average percent composition of CD3+ T cells per lymph node (50 ± 17%) and the average number of cells per 300-μm-thick lymph node slice ($(0.56 \pm 0.16) \times 10^6$ cells).[7] Assuming each murine lymph node slice had two T cell zones (most have one or two zones; see Figure S2), this yielded a rough order-of-magnitude estimate of 140,000 T cells per T cell zone in a lymph node slice.

As an example, using these numbers, for a T cell zone in which the oxygen consumption rate was $19 \times 10^{-4}$ mM/sec as measured by the optical assay, we estimated that $2.4 \times 10^{-4}$ pmol/min/cell was consumed.
